# Artificial light at nigh drives early dawn chorus onset times of the Saffron Finch (*Sicalis flaveola*) in an Andean city

**DOI:** 10.1101/2020.06.11.146316

**Authors:** Oscar Humberto Marín Gómez

**Affiliations:** Red de Ambiente y Sustentabilidad, Instituto de Ecología, A.C., Carretera antigua a Coatepec 351 El Haya, Xalapa 91070, Veracruz, Mexico

**Keywords:** Andean cities, Neotropics, song timing, singing routines, urbanization

## Abstract

Urban birds around the world have to cope with dominant city stressors as anthropogenic noise and artificial light at night by adjusting the temporal and spectral traits of their acoustic signals. It is widely known that higher anthropogenic noise and artificial light levels can disrupt the morning singing routines, but its influence in tropical urban birds remains poorly explored. Here, I assessed the association between light and noise pollution with the dawn chorus onset of the Saffron Finch (*Sicalis flaveola*) in an Andean city of Colombia. I studied 32 urban sites distributed in the north of the city, which comprise different conditions of urban development based on the built cover. I annotated the time when the first individual of the Saffron Finch was heard at each site and then I obtained anthropogenic noise and artificial light at night measurements using a smartphone. Findings of this study show that Saffron Finches living in highly developed sites sang earlier at dawn than those occupying less urbanized sites. Unexpectedly this timing difference was related to artificial lighting instead of anthropogenic noise, suggesting that artificial light could drive earlier dawn chorus in a tropical urban bird. Saffron Finches could take advantage of earlier singing for signaling territorial ownership among neighbors, as expected by the social dynamic hypothesis. However, findings of this study should be interpreted carefully because the dawn chorus is a complex phenomenon influenced by many multiple factors. Future studies need to assess the influence of ALAN on the dawn chorus timing of Neotropical urban birds by taking into account the influence of confounding factors related to urbanization as well as meteorological, ecological, and social drivers.

## INTRODUCTION

The daily singing routines of birds typically involve two peaks of high vocal activity performed by different individuals of multiple species around dawn and dusk (Leopold & Eynon 1961, Staicer et al. 1996, Catchpole & Slater 2008). Those choruses are a dominant component of the biophony of terrestrial soundscapes (Farina & Ceraulo 2017) and represent an intriguing feature of the avian natural history (Staicer et al. 1996). In particularly, avian dawn choruses have received important attention due to their consequences for fitness (Staicer et al. 1996, Gil & Lluisa 2020). Given the complexity of this phenomenon, multiple hypotheses have been proposed to explain why birds sing more at dawn than during daytime, including physiological (circadian rhythms on testosterone and melatonin); environmental (predation risk, acoustic transmission, inefficient foraging, body condition); and social factors (intrasexual and intersexual communication; see Staicer et al. 1996, Gil & Llusia 2020).

Urban environments have become important scenarios to understand how birds can deal and adapt to novel conditions generated by the environmental and ecological changes that urban sprawl implies (Gil & Brumm 2014). On this line, avian communication in cities have received much attention in recent years, providing evidence of acoustic adjustments to urbanization (reviewed by Slabbekoorn 2013). In general, birds well-adapted to urbanization have to cope with dominant urban stressors as anthropogenic noise and artificial light by adjusting the temporal and spectral traits of their acoustic phenotype (Slabbekoorn 2013, Gil & Brumm 2014). As noise levels peak at rush hours (mainly traffic and pedestrian activity) having earlier singing routines is a good strategy to avoid acoustic masking (Bergen & Abs 1997, Warren et al. 2006, Fuller et al. 2007, Gil et al. 2015, Dorado-Correa et al. 2016). Adjustments on structural traits as increasing minimum acoustic frequencies as well the amplitude of the vocalizations are another strategy to reduce acoustic masking and increase the active space for signaling (Brumm 2004, Gil & Brumm 2014, Sierro et al. 2017, Bermúdez-Cuamatzin et al. 2018).

Artificial light at night (ALAN) comprises any source of anthropogenic illumination at night, both indoor and outdoor, which drives changes on the physiology and behavior of the urban wildlife (Da Silva & Kempenaers 2017, Hopkins et al. 2018, Ouyang et al. 2018, Spoelstra et al. 2018). Studies from Palearctic and Nearctic cities suggest that ALAN disrupts the daily rhythms in songbirds by altering the circadian rhythms of sleep and hormone production (Dominoni 2015, De Jong et al. 2016, Hopkins et al. 2018, Russart & Nelson 2018). As ALAN generates similar light levels as those observed during natural twilight periods (i.e., transition between day and night when the sun drops under the horizon) circadian rhythms could be altered by the extension of twilight period (Secondi et al. 2020). For instance, some bird species commonly found in urban environments can extend their foraging and vocal activity to night-time in areas with high levels of light pollution (Fuller et al. 2007, MacGregor-Fors et al. 2011, Russ et al 2015, Leveau 2020). Additionally, diurnal birds exposed to ALAN tend to anticipate the onset of their dawn chorus (Bergen & Abs 1997, Miller 2006, Kempenaers et al. 2010). However, the relationship between noise and light pollution on singing routines such as dawn chorusing remains poorly understood in Neotropical urban birds (Dorado-Correa et al. 2016, Marín-Gómez & MacGregor-Fors 2019, Marín-Gómez et al. 2020).

In a pioneer study in the city of Bogotá, Dorado-Correa et al. (2016) assessed the influence of traffic noise and light pollutionon the singing behavior of the Rufous-collared Sparrow (*Zonotrichia capensis)*, a widely distributed species and a frequently dweller of some South American cities (Rising & Jaramillo 2020a). They found that light pollution was not related to earlier singing behavior, but Rufous-collared sparrows living in noisy urban sites showed earlier singing as a strategy to avoid acoustic masking by traffic later in the morning (Dorado-Correa et al. 2016). Another study in the city of Xalapa (Mexico), suggested that anthropogenic noise was related with earlier dawn singing onset and chorus peak in urban areas instead of light pollution (Marín-Gómez & MacGregor-Fors 2019). Given the recent interest on the study of the potential influence of both anthropogenic noise and ALAN with daily singing routines of tropical birds, in the present study I assessed the association between light and noise pollution with dawn chorus onset of the Saffron Finch (*Sicalis flaveola*) in different urbanization conditions of an Andean city in Colombia.

## Material and Methods

### Study species

The Saffron Finch is a small granivorous bird widely spread across lowlands (usually below 1000 m) of South America from Colombia to Argentina (Hilty et al. 2003). This finch inhabits pastures, semi-open areas with scattered bushes, lawns, and gardens in rural and urban areas (Hilty et al. 2003, Rising & Jaramillo 2020b). It is a secondary cavity nester that also uses abandoned nests from other species (Espinosa et al. 2017), artificial cavities including light poles and roofs in urban areas (Marín-Gómez *obs. pers*). Widely kept and popular as a cage-bird, this finch is very tolerant to human presence and a frequent visitor in urban feeders and gardens (Hilty et al. 2003, Rising & Jaramillo 2020b). Despite being a very common species, its ecology and behavior remain poorly studied (Benítez & Massonia 2018a, 2018b, Rising & Jaramillo 2020b). Saffron Finch males often sing exposed from a conspicuous perch (branch, post, power-line) and exhibited a very large and highly variable repertoire (25 ± 1.9 syllables), with short songs (2.1 ± 0.31 s) emitted at wide-frequency range (5.97 ± 0.12 kHz; Benítez & Massonia 2018a).

### Study sites and sampling design

I conducted this study in the north of Armenia, the capital of the Quindío Department, a city of 115 km^2^ with a population of 372,344 people. The city is located in the central Andes of Colombia at 1350-1550 m a.s.l., with an annual mean precipitation of 2,163 mm, a mean temperature of 21.8°C, and relative humidity ranging between 76 and 81% (Marín-Gómez et al. 2016). Currently, it is characterized by large buildings that contrast with small houses and green areas, urban parks, and corridors of remnant vegetation through the urban area.

I studied 32 urban sites (Figure 1) distributed in the north of the city which comprises different conditions of urban development from sparsely areas, typically green areas and urban parks, to highly developed sites as residential and commercial areas exposed to relatively high traffic noise and light pollution levels. Sampling sites were selected from a citywide study (designed to survey the taxonomical and functional bird diversity) and where Saffron finches was previously recorded (Marín-Gómez *obs. pers*). To ensure independence, sampling sites were spaced by a minimum distance of 300 m. These sites represented different ecological conditions of the urbanization across the city, defined by the percentage of built cover within a 50 m radius buffer for each sampling site measured from Google Maps.

**Figure 1.**
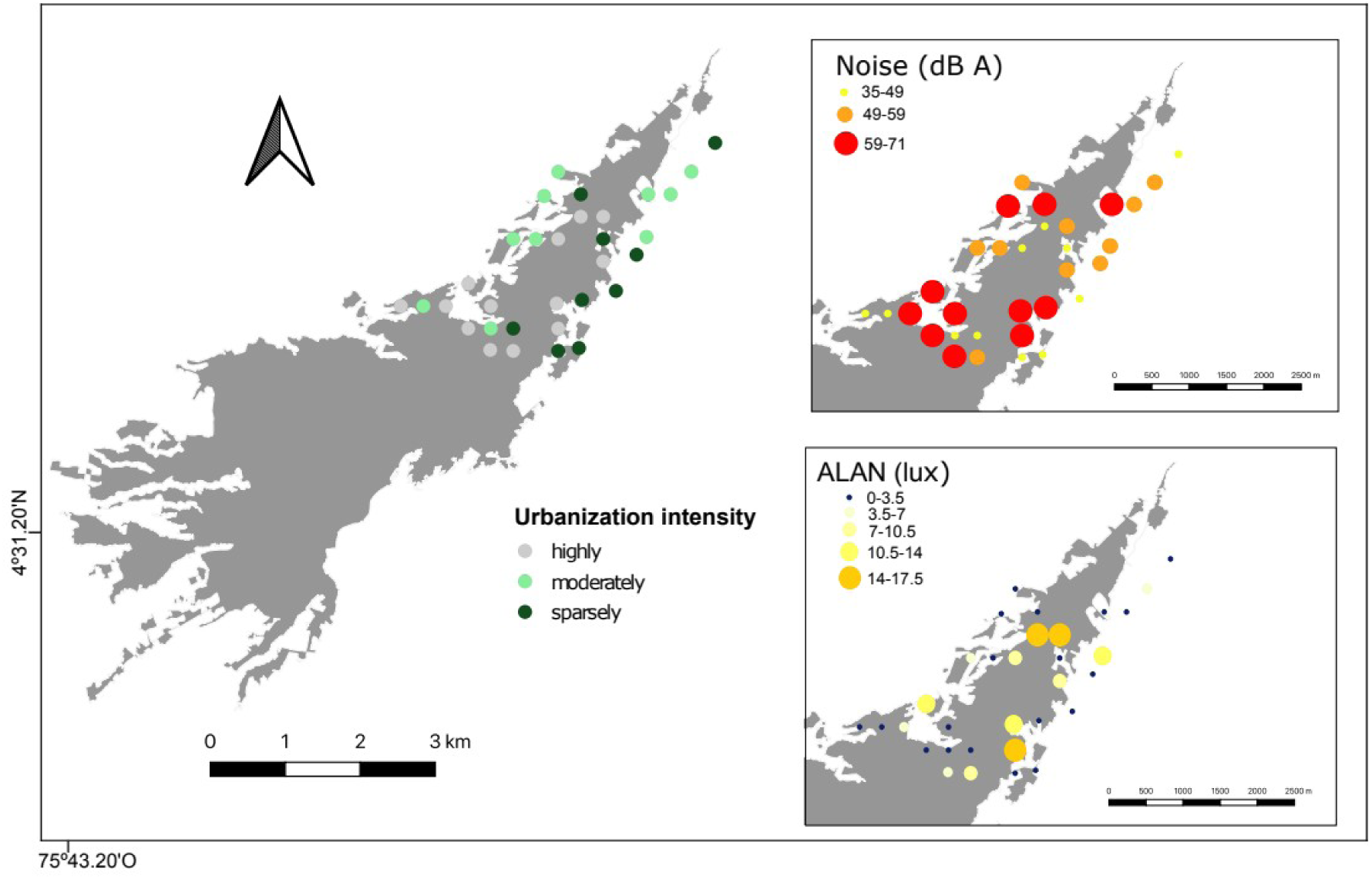
Map of the studied locations according to urbanization intensity, anthropogenic noise and artificial light levels.

Afterward, I classified each site based on the urban intensity as follows (MacGregor-Fors 2011): sparsely developed (0–33% built cover), moderately developed (34–66% built cover), and highly developed (67–100% built cover).

### Dawn chorus data and site variables

I visited each site once from Dec 2016 to Jan 2017 to record the time when the first individual of the Saffron Finch was heard (see Figure 2). Given that some songbirds can vocalize at night (Fuller 2007), I started the observations at nautical twilight of each sampling date (retrieved from the US Naval Observatory http://aa.usno.navy.mil). Then, I located the perch where each bird was first heard in order to record the maximum levels of both anthropogenic noise and artificial light at night. I measured anthropogenic noise and artificial light pollution levels using two smartphone apps: Lux Meter (Crunchy ByteBox) and Sound Meter app (Abc Apps) for ASUS Zenfone 2 smartphone. For this, I held the smartphone at 1.2 m and moved it to each cardinal point to record during a 1–min the maximum value of noise (dB) and then during a 1–min the maximum value of light pollution (lux). Although those measurements can not reflect real values as those obtained by sound meter and lux meter instruments, the results obtained could be useful as a proxy to describe noise and light pollution levels of a given environment (Kardous & Shaw 2014, Gutierrez-Martinez et al. 2017).

**Figure 2.**
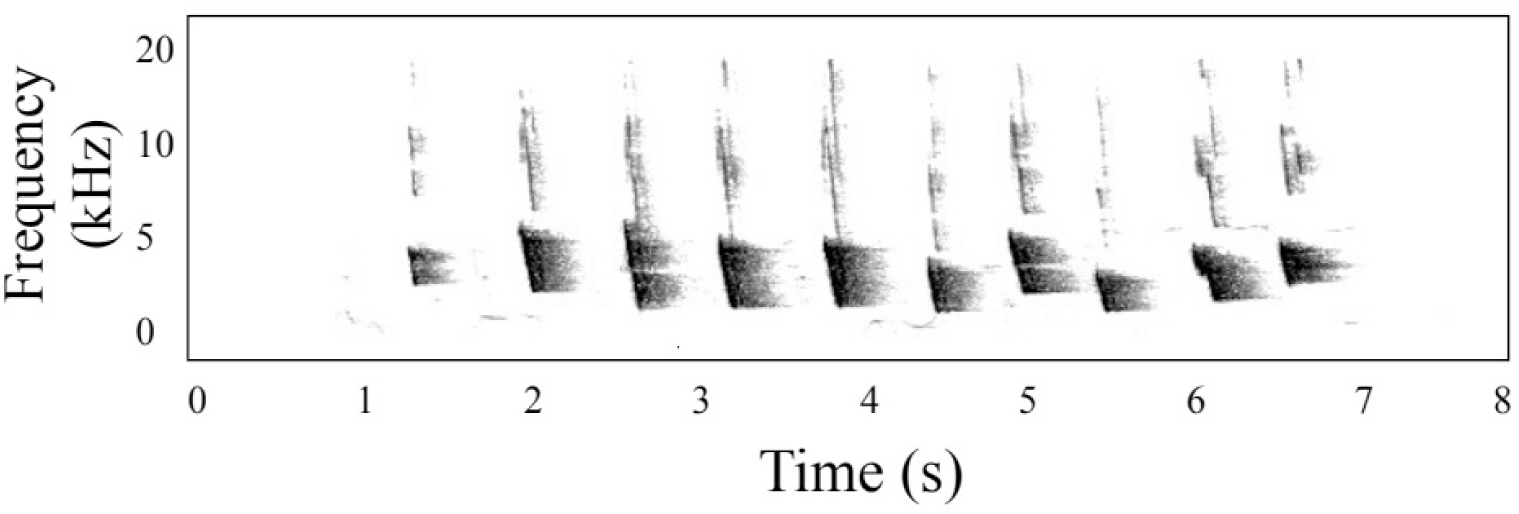
Spectrogram of the dawn song of the Saffron Finch recorded in this study.

### Data analyses

I tested noise and light pollution levels variation across the urbanization conditions using Kruskal-Wallis test. Then, I assessed the correlation between both independent variables with Pearson correlation coefficient. I used linear models to test relationships between the onset of dawn chorus with ALAN and anthropogenic noise levels as well as the interactions between them. The dependent variable was the dawn chorus onset time calculated as the difference between the time of the first song of the Saffron Finch relative to the time of sunrise, where negative values represent onset times before sunrise. The independent variables were noise levels (dB(A)), and ALAN (lux). The model was validated through diagnostic plots of residual heterogeneity and homoscedasticity assumptions (Crawley 2012). All statistical analyses were carried out using R 3.4.

## Results

ALAN and anthropogenic noise levels varied across sampling sites (ALAN: H_2,32_= 16.0, p < 0.01; noise: H_2,32_= 5.6, p = 0.06). Highly urbanized sites showed higher levels of both ALAN and anthropogenic noise than sparsely developed sites (Table 1). Interestingly, ALAN and anthropogenic noise were not correlated (r = 0.24; N=32, p= 0.09) suggesting absence of collinearity between both variables. The Saffron Finch was typically a dawn singer which chorus onset times varied from 51 min to 10 min before sunset (−34.0 ± 12.4 min) and occurred earlier in highly urbanized sites (Figure 3). Saffron Finches started to sing earlier in sites exposed to higher ALAN and anthropogenic noise levels (Figure 4). However, only ALAN predicted chorus onset (Table 2), suggesting that birds living in highly illuminated sites (> 7.5 lux, onset: -47.2 ± 3.3 min) started to sing 20 min earlier than birds exposed to low illuminated areas (< 7.5 lux, onset: -28.8 ± 10.7 min).

**Table 1.** Variation of anthropogenic noise and artificial light levels among the studied urban conditions in the city of Armenia, Colombia

| Urbanization condition | Descriptive statistic | Artificial light (lux) | Noise levels at chorus onset (dB(A)) |
| --- | --- | --- | --- |
| Highly developed (n = 13) | min | 0.5 | 35 |
|  | max | 17.5 | 70 |
|  | average (SD) | 8.85 ± 5.58 | 55.54 ± 10.88 |
| Moderately developed (n = 10) | min | 0.5 | 44.5 |
|  | max | 13 | 71 |
|  | average (SD) | 3.55 ± 3.66 | 54.06 ± 9.69 |
| Sparsely developed (n = 9) | min | 0 | 37 |
|  | max | 3.5 | 59.5 |
|  | average (SD) | 0.83 ± 1.17 | 46.16 ± 8.32 |

**Table 2.** Results of a linear model testing association among the dawn chorus onset of *Sicalis flaveola* with anthropogenic noise and artificial light (ALAN) in the city of Armenia, Colombia.

| Predictors | Estimate | Std error | t-value | p-value |
| --- | --- | --- | --- | --- |
| Intercept | -6.27 | 0.75 | -0.58 | 0.564 |
| Noise | -0.38 | 0.21 | -1.83 | 0.077 |
| ALAN | -3.76 | 1.72 | -2.19 | 0.037 |
| Noise:ALAN | 0.04 | 0.03 | 1.27 | 0.212 |

**Figure 3.**
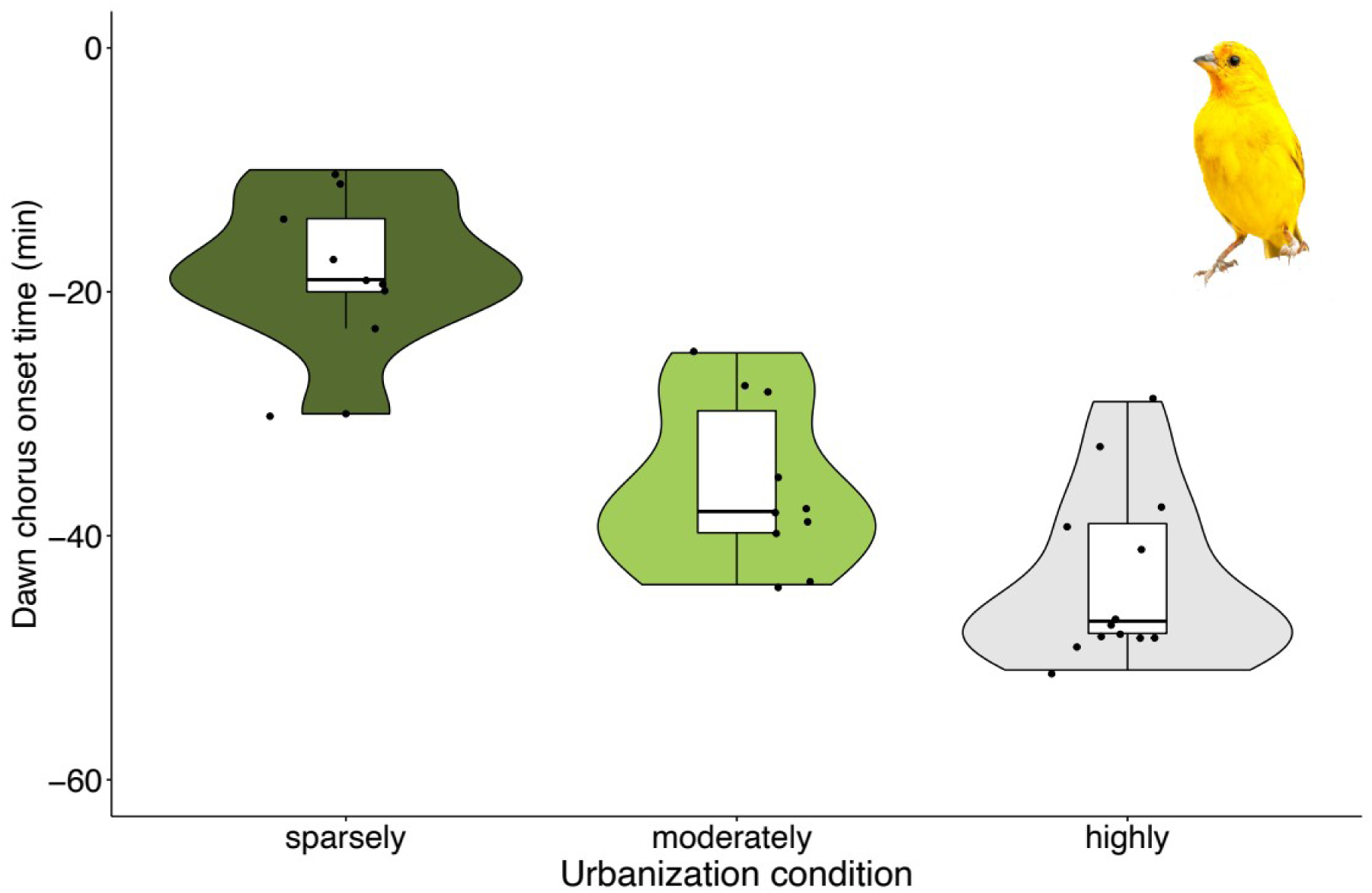
Violin plot showing the variation of dawn chorus onset times of the Saffron finch for each urbanization condition in the city of Armenia, Colombia. Sparsely (dark green), moderately (light green), and highly developed (gray).

**Figure 4.**
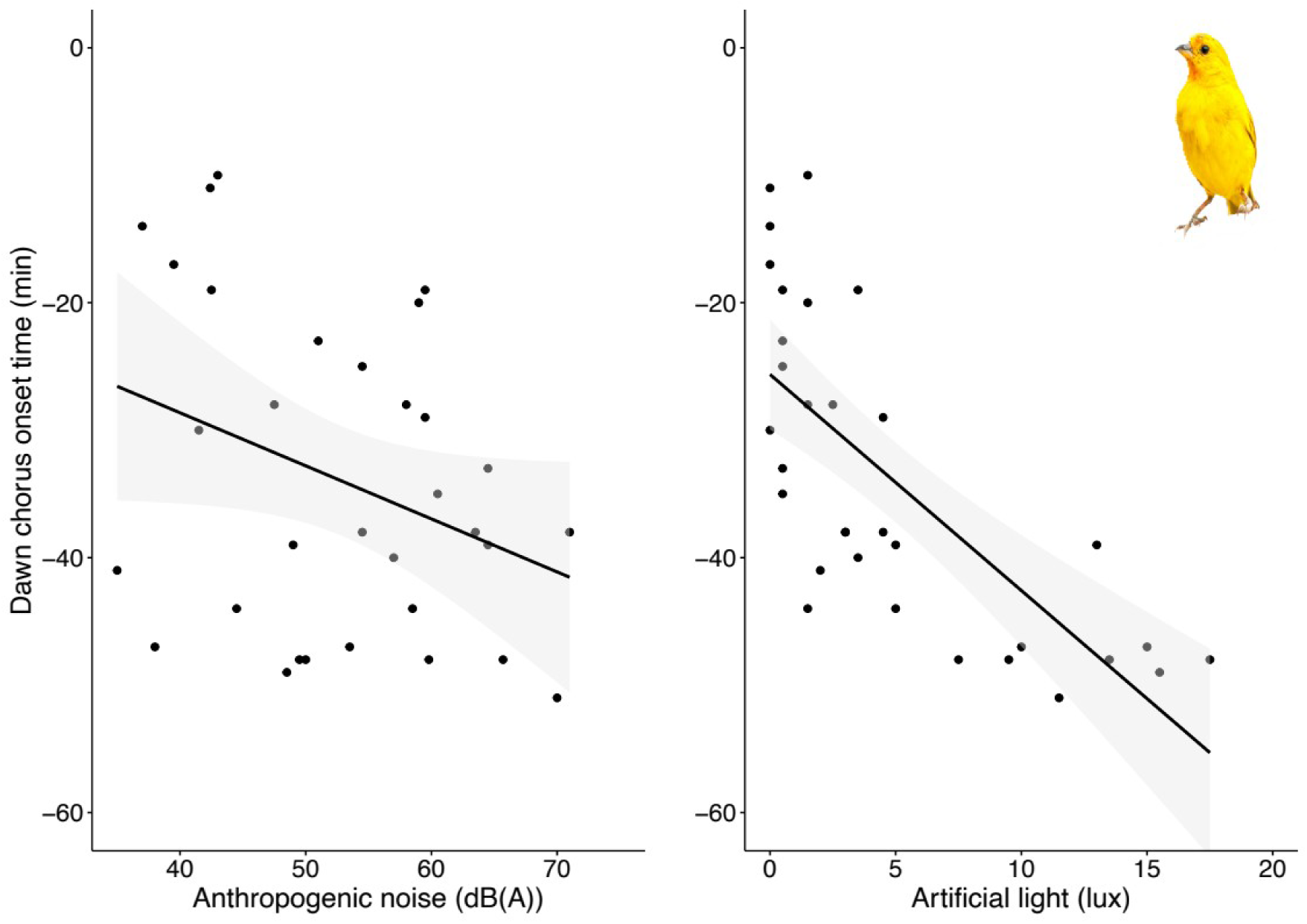
Relationships among the dawn chorus onset of the Saffron finch with built cover, anthropogenic noise, and ALAN. Grey bands represent 95% confidence intervals

## Discussion

Saffron finches started their activity earlier in highly urbanized sites as previously found for other songbirds (Bergen & Abs 1997, Fuller et al. 2007; Arroyo-Solís et al. 2013, Da Silva et al. 2015, Dorado-Correa et al. 2016, Da Silva & Kempenaers 2017). However, earlier chorus onset was not related to anthropogenic noise levels at the moment of the singing, an unexpected result given the growing body of evidence supporting anthropogenic noise as the main factor driving the timing of singing behavior in urban birds in temperate and even tropical species (Gil et al. 2015, Dorado-Correa et al. 2016, Sierro et al. 2017, Marín-Gómez & MacGregor-Fors 2019). Having found no association between chorus onset and noise levels does not imply that Saffron finches are not affected by noise. Noise levels at rush hours (around sunrise and early morning) in highly urbanized areas tend to be higher than at dawn (Gil et al. 2015, Dominoni et al. 2016, Sierro et al. 2017). Unfortunately, data on noise variation from dawn to morning are unknown to test whether Saffron finches sing earlier to avoid acoustic masking at rush hours. Additionally, the start of singing behavior is only one component of the singing routines of birds (Naguib et al 2019). The time in which a bird broadcasts more vocalizations (chorus peak) could be also important. For instance, a recent study in a neotropical city shows no relationship between noise levels and chorus onset but chorus peak (Marín-Gómez & MacGregor-Fors 2019).

Given that the length of photoperiod tends to be more constant towards the Equator (Secondi et al. 2020), the annual reproductive cycle of most tropical birds may be less affected by variations in day length than temperate species (Stutchbury & Morton 2008, Stouffer et al. 2013, but see Hau 2001, Goymann et al. 2012). Therefore, it is expected that the breeding phenology of tropical bird species could be less affected by ALAN (i.e., Da Silva & Kempenaers 2017). For instance, recent studies suggest that anthropogenic noise but not ALAN is related to the timing of singing routines in neotropical urban birds (Dorado-Correa et al. 2016, Marín-Gómez & MacGregor-Fors 2019, Marín-Gómez et al. 2020). However, findings of this study show, for the first time, that high levels of ALAN were related to the earlier dawn chorus in a tropical species. Some explanations could be proposed to explain this pattern. First, some tropical birds can be sensitive to slight photoperiodic variations and use it to initiate reproductive activity (Hau et al. 1998, Hau 2001, Goymann et al. 2012), then Saffron finches could detect variations on natural light and ALAN. Second, birds living in urban sites exposed to different levels of ALAN (0.05–6 lux) can wake up earlier and sleeping less (De Jong et al. 2016, Raap et al. 2017, Ouyang et al. 2018), therefore Saffron Finches could anticipate the onset of their daily activities by waking up earlier and then start to sing. In fact, in the studied city Saffron Finches as well as other common species as the Vermilion Flycatcher (*Pyrocephalus rubinus*) and the Tropical Kingbird (*Tyrannus melancholicus*) show nocturnal singing near to lamp posts (Marín-Gómez *obs. pers*).

Earlier activities driven by ALAN can affect predator-prey interactions, as well as breeding success (Miller 2006, Kempenaers et al 2010). Studies in temperate cities indicate that an increase of apparent day length as a result of ALAN may provide more time for foraging but also for social signaling (Ouyang et al. 2018). Earlier activities of Saffron Finches in highly illuminated areas could not be related to improving foraging times because as seed-eaters this species cannot forage for insects attracted to light poles as typically do some tropical flycatchers (MacGregor-Fors et al. 2011, Marín-Gómez obs. pers). Therefore, Saffron Finches could take advantage of earlier singing for signaling territorial ownership among neighbors, as expected by the social dynamic hypothesis (Staicer et al. 1996, Gil & Llusia 2020). However, results of this study should be interpreted carefully because the dawn chorus is a multifactorial phenomenon influenced by many environmental factors like noise, light intensity, cloud cover, moonlight, temperature, and food supply; as well biological factors as breeding season, home range size, eye-size, foraging guild, hetero-specific neighbors occurrence, and population density of conspecific neighbors (Staicer et al. 1996, Malavasi & Farina 2013, Da Silva et al. 2017, Gil & Llusia 2020). Previous studies have tested the influence of environmental factors on dawn chorus timing in tropical birds without considering biological factors (Dorado-Correa et al. 2016, Marín-Gómez & MacGregor-Fors 2019, Marín-Gómez et al. 2020). Moreover, measurements of noise and light pollution presented here are biased because they were based on smartphone applications instead of precise equipment as sound and light meters. Further studies need to obtain precise measurements of ALAN and noise levels variation at both daytime and nighttime (e.g., Marín-Gómez et al. 2020) in order to assess the influence of urban-related factors (noise, ALAN, temperature) on daily singing routines in tropical birds.

In summary, findings of this study show that Saffron finches living in highly developed sites of an Andean city sang earlier at dawn than those occupying less urbanized sites. Unexpectedly this timing difference was related to artificial lighting instead of anthropogenic noise, suggesting that artificial light could drive earlier dawn chorus in a tropical urban bird. Further studies needs to take into account the influence of multiple potential drivers as meteorological (cloudiness, temperature, moon-light), ecological (body condition, food supply, predation risk), and social factors (density of neighbors) on daily singing routines in neotropical urban birds.

## ACKNOWLEDGEMENTS

To Margarita Lopez Garcia for helping in the field work and performing the map and to Ina Susana Falfán for providing shapefiles. I was supported by the scholarship provided by the National Council of Science and Technology (CONACYT 417094), and the Doctoral Program of the Instituto de Ecología, A.C. (INECOL).

## REFERENCES

Arroyo-Solís, A, JM Castillo, E Figueroa, JL López-Sánchez & H Slabbekoorn (2013) Experimental evidence for an impact of anthropogenic noise on dawn chorus timing in urban birds. Journal of Avian Biology 44: 288–296.

Benítez Saldívar, M. & V Massoni (2018a) Song structure and syllable and song repertoires of the Saffron Finch (*Sicalis flaveola pelzelni*) breeding in Argentinean pampas. Bioacoustics 27: 327–340.

Benítez Saldívar, M. & V Massoni (2018b) Lack of conspecific visual discrimination between second-year males and females in the Saffron Finch. PloS one 13: e0209549.

Bergen, F & M Abs (1997) Verhaltensökologische Studie zur Gesangsaktivität von Blaumeise (*Parus caeruleus*), Kohlmeise (*Parus major*) und Buchfink (*Fringilla coelebs*) in einer Großstadt. Journal für Ornithologie 138: 451–467.

Bermúdez-Cuamatzin, E, M López-Hernández, J Campbell, I Zuria & H Slabbekoorn (2018) The role of singing style in song adjustments to fluctuating sound conditions: A comparative study on Mexican birds. Behavioural Processes 157: 645–655.

Brumm, H (2004) The impact of environmental noise on song amplitude in a territorial bird. Journal of Animal Ecology 73: 434–440.

Crawley, MJ (2012) The R book. John Wiley & Sons.

Da Silva, A, M Valcu & B Kempenaers (2015) Light pollution alters the phenology of dawn and dusk singing in common European songbirds. Philosophical Transactions of the Royal Society B: Biological Sciences 370: 20140126–20140126.

Da Silva, A & B Kempenaers (2017) Singing from North to South: Latitudinal variation in timing of dawn singing under natural and artificial light conditions. Journal of Animal Ecology 86: 1286–1297.

De Jong, M, L Jeninga, JQ Ouyang, K van Oers, K Spoelstra & M Visser (2016) Dose-dependent responses of avian daily rhythms to artificial light at night. Physiology & Behavior 155: 172–179.

Dominoni, DM (2015) The effects of light pollution on biological rhythms of birds: an integrated, mechanistic perspective. Journal of Ornithology 156(1), 409–418.

Dorado-Correa, AM, M Rodríguez-Rocha & H Brumm (2016) Anthropogenic noise, but not artificial light levels predicts song behaviour in an equatorial bird. Royal Society Open Science 3: 160231.

Espinosa, C, L Cruz-Bernate & G Barreto (2017) Reproductive biology of *Sicalis flaveola* (Aves: Thraupidae) in Cali, Colombia. Boletín Científico Centro de Museos Museo de Historia Natural 21: 101–114.

Farina, A, M Ceraulo, C Bobryk, N Pieretti, E Quinci & E Lattanzi (2015) Spatial and temporal variation of bird dawn chorus and successive acoustic morning activity in a Mediterranean landscape. Bioacoustics 24: 269–288.

Fuller, RA, PH Warren & K Gaston (2007) Daytime noise predicts nocturnal singing in urban robins. Biology Letters 3: 368–370.

Gil, D & H Brumm (2014) Avian urban ecology. OUP Oxford.

Gil, D, M Honarmand, J Pascual, E Pérez-Mena & C Macías García (2015) Birds living near airports advance their dawn chorus and reduce overlap with aircraft noise. Behavioral Ecology 26: 435–443.

Gil, D & D Llusia (2020) The bird dawn chorus revisited. In Coding strategies in vertebrate acoustic communication (pp. 45–90). Springer, Cham.

Goymann, W, B Helm, W Jensen, I Schwabl & IT Moore (2012) A tropical bird can use the equatorial change in sunrise and sunset times to synchronize its circannual clock. Proceedings of the Royal Society B: Biological Sciences 279: 3527–3534.

Gutierrez-Martinez, JM, A Castillo-Martinez, JA Medina-Merodio, J Aguado-Delgado & JJ Martinez-Herraiz (2017) Smartphones as a light measurement tool: Case of study. Applied Sciences 7: 616.

Hau, M, M Wikelski M & JC Wingfield (1998) A Neotropical forest bird can measure the slight changes in tropical photoperiod. Proceedings of the Royal Society of London Series B: Biological Sciences 265: 89–95.

Hau, M (2001) Timing of breeding in variable environments: tropical birds as model systems. Hormones and Behavior 40: 281–290.

Hilty, SL, G Tudor & JA Gwynne (2003). Birds of Venezuela. Princeton University Press.

Hopkins, GR, KJ Gaston, ME Visser, MA Elgar & TM Jones (2018) Artificial light at night as a driver of evolution across urban–rural landscapes. Frontiers in Ecology and the Environment, 16(8): 472–479.

Kardous, C. & PB Shaw (2014) Evaluation of smartphone sound measurement applications. The Journal of the Acoustical Society of America 135: EL186–EL192.

Kempenaers, B, P Borgström, P Loës, E Schlicht & M Valcu (2010) Artificial night lighting affects dawn song, extra-pair siring success, and lay date in songbirds. Current Biology 20: 1735–1739.

Leopold, A & AE Eynon (1961) Avian daybreak and evening song in relation to time and light intensity. The Condor 63: 269–293.

Leveau, LM (2020) Artificial light at night (ALAN) is the main driver of nocturnal feral pigeon (*Columba livia f. domestica*) foraging in urban areas. Animals 10: 554.

MacGregor-Fors, I (2011). Misconceptions or misunderstandings? On the standardization of basic terms and definitions in urban ecology. Landscape and Urban Planning 100: 347–349.

MacGregor-Fors, I, A Blanco-García, C Chávez-Zichinelli, E Maya-Elisararrás, L Mirón, L Morales-Pérez & JE Schondube (2011) Relación entre la presencia de luz artificial nocturna y la actividad del mosquero cardinal (*Pyrocephalus rubinus*). El Canto del Centzontle 2: 64–71.

Malavasi, R & A Farina (2013) Neighbours’ talk: interspecific choruses among songbirds. Bioacoustics 22: 33–48.

Marín-Gómez, OH, JI Garzón Zuluaga, DM Santa-Aristizabal, J Hernán López & MM López-García (2016) Use of urban areas by two emblematic and threatened birds in the central Andes of Colombia. Revista Brasileira de Ornitologia 24: 260–266.

Marín-Gómez, O. & I MacGregor-Fors (2019). How early do birds start chirping? dawn chorus onset and peak times in a Neotropical city. Ardeola 66: 327–341.

Marín-Gómez, OH,M García-Arroyo, CE Sánchez-Sarria, JR Sosa-López, D Santiago-Alarcon & I MacGregor-Fors (2020) Nightlife in the city: drivers of the occurrence and vocal activity of a tropical owl. Avian Research 11: 1–14.

Miller, MW (2006) Apparent effects of light pollution on singing behavior of American Robins. The Condor 108: 130–139.

Naguib, M, J Diehl, K Van Oers & L Snijders (2019) Repeatability of signalling traits in the avian dawn chorus. Frontiers in Zoology 16: 27.

Ouyang, JQ, S Davies & D Dominoni (2018) Hormonally mediated effects of artificial light at night on behavior and fitness: linking endocrine mechanisms with function. Journal of Experimental Biology 221: jeb156893.

Raap, T, J Sun, R Pinxten & M Eens (2017) Disruptive effects of light pollution on sleep in free-living birds: Season and/or light intensity-dependent?. Behavioural Processes 144: 13–19.

Rising, J. & A Jaramillo (2020a) Rufous-collared Sparrow (Zonotrichia capensis), version 1.0. In Birds of the World (Del Hoyo, J, A Elliott, J D Sargatal, A Christie & E de Juana (Editors). Cornell Lab of Ornithology, Ithaca, NY, USA. https://doi.org/10.2173/bow.rucspa1.01

Rising, J. & A Jaramillo (2020b) Saffron Finch (*Sicalis flaveola*), version 1.0. In Birds of the World (Del Hoyo, J, A Elliott, J D Sargatal, A Christie & E de Juana (Editors). Cornell Lab of Ornithology, Ithaca, NY, USA. https://doi.org/10.2173/bow.saffin.01

Russ, A, A Rüger & R Klenke (2015) Seize the night: European Blackbirds (*Turdus merula*) extend their foraging activity under artificial illumination. Journal of Ornithology 156: 123–131.

Russart, K. & RJ Nelson (2018) Artificial light at night alters behavior in laboratory and wild animals. Journal of Experimental Zoology Part A: Ecological and Integrative Physiology 329: 401–408.

Secondi, J, A Davranche, M Théry, N Mondy & T Lengagne (2020). Assessing the effects of artificial light at night on biodiversity across latitude–Current knowledge gaps. Global Ecology and Biogeography 29: 404–419.

Sierro, J, E Schloesing, I Pavón & D Gil (2017) European Blackbirds exposed to aircraft noise advance their chorus, modify their song and spend more time singing. Frontiers in Ecology and Evolution 5: 68.

Slabbekoorn, H (2013) Songs of the city: noise-dependent spectral plasticity in the acoustic phenotype of urban birds. Animal Behaviour 85: 1089–1099.

Spoelstra, K, I Verhagen, D Meijer & ME Visser (2018) Artificial light at night shifts daily activity patterns but not the internal clock in the great tit (*Parus major*). Proceedings of the Royal Society of London Series B: Biological Sciences 285: 20172751.

Staicer, C, D Spector & A Horn (1996) The dawn chorus and other diel patterns in acoustic signalling. In Kroodsma, DE & EH Miller (eds.), Ecology and evolution of acoustic communication in birds, pp. 426–453. Cornell University Press., Ithaca, New York.

Stouffer, PC, EI Johnson & RO Bierregaard Jr (2013) Breeding seasonality in central Amazonian rainforest birds. The Auk 130: 529–540.

Stutchbury, B. & E Morton, ES (2008) Recent advances in the behavioral ecology of tropical birds: the 2005 Margaret Morse Nice Lecture. The Wilson Journal of Ornithology 120: 26–37.

Warren, PS, M Katti, M Ermann & A Brazel (2006) Urban bioacoustics: it’s not just noise. Animal Behaviour 71: 491–502.

